# Thylakoid plasticity primes photoprotection in diatoms

**DOI:** 10.64898/2026.10.27.761787

**Authors:** David Perez Compte, Dimitri Tolleter, Clarisse Uwizeye, Cécile Giustini, Lydia Pernet, Daphna Fenel, Christine Moriscot, Wojciech Wietrzynski, Renáta Ünnep, Gergely Nagy, Győző Garab, Pierre-Henri Jouneau, Guy Schoehn, Anja Krieger-Liszkay, Benjamin D. Engel, Pascal Albanese, Giovanni Finazzi, Serena Flori

**Affiliations:** Laboratoire de Physiologie Cellulaire et Végétale, Université Grenoble Alpes, CNRS, CEA, INRAE, IRIG-DBSCI, 17 rue des Martyrs, 38000 Grenoble, France; Institut de Biologie Structurale (IBS), 71 Av. des Martyrs, 38000 Grenoble, France; Biozentrum, University of Basel, Spitalstrasse 41, 4056 Basel, Switzerland; Budapest Neutron Centre, HUN-REN Centre for Energy Research, Institute for Energy Security and Environmental Safety, Hungary H-1121 Budapest Konkoly-Thege Miklós út 29-33; Laboratory for Neutron Scattering and Imaging, Paul Scherrer Institute, Villigen, Switzerland; Neutron Scattering Division, Oak Ridge National Laboratory, Oak Ridge, Tennessee, USA; Institute of Plant Biology, HUN-REN Biological Research Centre, Szeged, Temesvári körút. 62, 6726 Hungary; Department of Physics, Faculty of Science, University of Ostrava, Chittussiho 10, 702 00 Ostrava, Czech Republic; Université Grenoble Alpes, CEA, IRIG-MEM, Grenoble, France; Université Paris-Saclay, Institute for Integrative Biology of the Cell (I2BC), CEA, CNRS, 91190 Gif-sur-Yvette, France; EDyP, U1292, INSERM/CEA/UGA/IRIG, 17 rue des Martyrs, 38000 Grenoble, France

## Abstract

For many millions of years, diatoms have been major primary producers in marine ecosystems, supporting food webs and contributing to carbon sequestration through the export of photosynthetically fixed carbon to the deep ocean, mitigating atmospheric carbon dioxide levels and global climate. A key factor in their ecological success is their ability to rapidly adapt to fluctuating light, balancing efficient light harvesting under dim conditions with energy dissipation under excess light. However, the mechanisms behind this regulation have not been fully elucidated. This study shows that several processes contribute to dissipating excess excitation energy in diatoms exposed to light stress: (i) the detachment of the oxygen-evolving complex proteins from the PSII core, leading to energy dissipation via charge recombination, (ii) Non-Photochemical Quenching (NPQ) in the antenna, i.e., thermal dissipation of excited states and (iii) excitation energy transfer from PSII to PSI via energy spillover. The latter process is observed in parallel with high light-induced topological changes of the thylakoids that may favour transient interactions between the photosystems. Along with the accumulation of light-stress-protecting carotenoids on longer time-scales, this ensemble of photoprotective mechanisms provides a dynamic and flexible response to the light challenges expected in coastal marine habitats.

## Introduction

Photosynthesis is an ancient process that converts light energy into chemical energy, fixing CO_2_ into organic matter and releasing O_2_ through water oxidation, across most environments on Earth. Aquatic photosynthesis accounts for a substantial fraction of global carbon fixation and primary productivity ^1^, and in modern oceans, diatoms are responsible for a major part of this productivity ^2^, thus being one of the most successful and cosmopolitan phytoplankton groups^3,4^. Several explanations have been proposed for this capacity to adapt to different environments, including the ability to continuously adjust light absorption and utilization^5,6^.

Diatoms contain a plastid of secondary endosymbiotic origin ^7,8^, which occupies approximately 30% of the entire cell volume ^6^. This plastid is bounded by a four-membrane envelope, a hallmark of its secondary endosymbiotic origin ^9,10^. Diatom photosynthetic membranes (thylakoids) host the same core components of the photosynthetic electron transport chain as *Viridiplantae*: Photosystem I and II (PSI and PSII), which perform the photochemical light conversion; the cytochrome *b*_6_*f* complex, connecting the two photosystems; and the ATP synthase, which couples the proton motive force to ATP production. Unlike the stacked grana and unstacked stroma lamellae of plants, the majority of thylakoids within diatom chloroplasts are organized in long triplet stacks of three thylakoids, connected by occasional single thylakoids ^10,11^. A common organizing principle of thylakoid stacking in plants ^12,13^, green algae ^14^, and diatoms ^5^ is lateral heterogeneity, which restricts PSII to appressed membrane domains and PSI to non-appressed membranes. However, the long thylakoid triplets in diatoms result in a different challenge for photosynthetic regulation compared to the membrane architecture found in *Viridiplantae*.

Diatoms also possess a sophisticated and unique machinery to collect light in their natural environment^15–17^, including specific antenna complexes functioning in the light harvesting or light dissipation modes. In the latter configuration, they dissipate excess absorbed energy in a process called Non-Photochemical Quenching (NPQ) ^15–17^. The NPQ capacity of diatoms^18,19^ can exceed the one observed in most phototrophs ^18–20^. Its levels are regulated by two parallel xanthophyll cycles ^21–23^ and by the accumulation of LHCX proteins ^24–26^. As in plants, NPQ in diatoms cannot be fully explained by a luminal-pH sensitive dissipation process, the so-called high dependent quenching (qE) ^18,25^. This finding suggests the presence of other protective mechanisms, raising the question of their origin and regulation.

The reversible association of antenna complexes with either PSI or PSII, known as state transitions, is a mechanism present in plants and green algae, but has not been observed in diatoms^27–29^. Instead, the occurrence of energy quenching inside the PSII reaction centre has been described ^18,30^. Finally, we highlight here the presence of another PSII-protective process, namely excitation energy spillover (hereafter ‘spillover’), in which PSII directly transfers energy to PSI upon excitation. This phenomenon appears when *Phaeodactylum tricornutum* cells are exposed to high light (HL), disappearing within minutes upon re-exposure to low light (LL) during the recovery phase. We interpret its occurrence in terms of a structural rearrangement of the photosynthetic membranes, lowering the segregation between the two photosystems and favouring their energy exchange. This remodelling of thylakoid membranes is reversible and is observed in diatoms upon HL exposure and subsequent relaxation to LL under physiological conditions.

Overall, non-photochemical quenching at the reaction center (RC-NPQ), and within the antennae (antenna-NPQ), and photochemical quenching by PSI through spillover act synergistically as complementary photoprotective mechanisms, enabling efficient energy dissipation in highly dynamic light environments.

## Results

### Phenotypic analysis of *P. tricornutum* cells reveals the existence of a ‘non-canonical’ NPQ process

Acclimation of diatom cells to different light intensities over several generations alters their photosynthetic properties, including antenna size, photochemical activity and structural dynamics of PSII (Fv/Fm) and NPQ ^31,32^. Subcellular changes in mitochondrial and plastid volumes and their interactions have been reported in cultures grown under different light intensities for several generations ^6^, ultimately affecting the photosynthetic and respiratory activities.

Here, we describe photoprotective responses occurring on a much shorter time scale, under conditions that mimic the light variations in the natural environment ^33^. We grew algae in low light (LL: 40 µmol photons m^-2^ s^-1^, cyan color) and then shifted them to higher light conditions (HL: 350 µmol photons m^-2^ s^-1^, purple color) for up to 2 hours. We examined their physiological responses and complemented them with biochemical and structural characterization. In HL, we observed a diminished photosynthetic activity (Fv/Fm, Fig. 1A) and photoprotection ability (NPQ, Fig. 1B). These effects were reversible upon reverting to the initial LL conditions (grey color). In parallel, we also observed an accumulation of the carotenoid pool affecting NPQ (diatoxanthin). This change was reversible in LL conditions (Fig. 1C). Finally, HL exposure led to a decrease in net oxygen evolution, which was also reversible upon switching to LL conditions (Fig. 1D).

**Figure 1.**
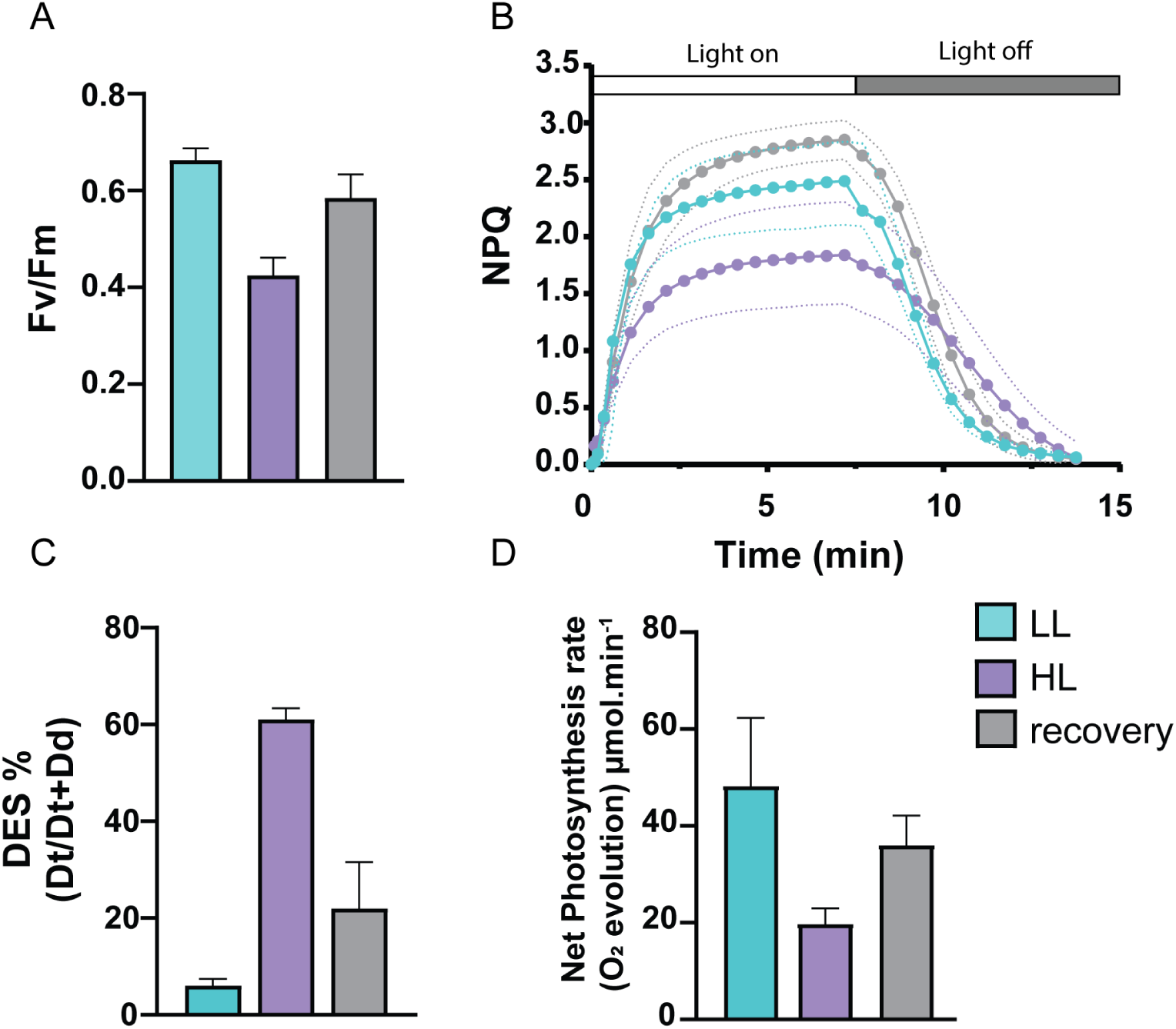
Photosynthetic properties and photoprotective responses of *P. tricornutum* cells under LL, HL and recovery conditions. (**A**) Maximum quantum yield of photosystem II (Fv/Fm) (**B**). NPQ kinetics over a 15-minute time course and standard deviation (in dotted lines). (**C**) De-epoxidation ratio of the xanthophyll cycle pigments, expressed as the ratio Dt/(Dt + Dd) (diadinoxanthin (Dd); diatoxanthin (Dt)). (**D**) Net photosynthesis rates (O_2_ evolution) measured at 298 µmol photons m⁻² s⁻¹ light intensity. Values represent mean ± standard deviation in at least two independent experiments (panels C see also Supplementary table1, and D) or more than three independent experiments (panels A and B).

Altogether, the data support the existence of a photoprotective response in diatoms, but reveal an apparent contradiction when compared to ‘textbook’ responses. Indeed, the accumulation of NPQ-effector carotenoids (Dt) that is observed upon light exposure does not enhance this process, as usually observed ^34^. Instead, NPQ decreased despite higher Dt levels, in parallel with diminished electron flow and PSII photochemical capacity. These physiological changes poorly correlate with modifications in the cell proteome: comparative proteomic analysis (LC-MS/MS) of whole-cell lysates from HL-versus LL-treated *P. tricornutum* revealed only a few significant changes among the 1790 identified proteins (adjusted p-value < 0.05 and |log_2_ FC| > 1; Fig. S1). Among the few differentially abundant proteins, the antenna subunits Lhcf15 and Lhcr6 were downregulated (Log₂FC = -1.18) and upregulated (Log₂FC = +2.47), respectively, after 2 h of HL exposure. These contrasting responses are consistent with their distinct roles in photoacclimation: Lhcf15 forms a putative red-shifted antenna associated with red-light acclimation ^35,36^, whereas Lhcr6 is induced by HL and has been previously implicated in photoprotection^31,37^.

These data suggest that the light acclimation responses above can neither be explained by a significant alteration of the cellular proteome, nor by a ‘classic’ NPQ induction sequence, in which HL exposure triggers xanthophyll de-epoxidation and NPQ protein induction/activation, ultimately leading to fluorescence quenching ^38^. Instead, they may be interpreted, at least to some extent, within the framework of a different type of photoprotective response.

### Biochemical and functional features of HL-exposed *P. tricornutum* cells

The RC-NPQ has been previously reported in diatoms ^18^ and displays functional characteristics similar to those of the reversible form of photoinhibition reported in earlier studies ^39–41^. We therefore tested whether the quenching characterised in Fig. 1 could reflect the same reversible photoinhibition state. We measured the fluorescence decay after a saturating single turnover flash. In the absence of DCMU, the curves could be fitted with two exponential decay functions (represented by the dotted lines, Fig. 2A). According to de Wijn and van Gorkom (2001) ^42^, these phases correspond to the electron transfer Q_A_^-^ to Q_B_ or Q_B_^-^ (fast phase), and to PQ binding to the empty Q_B_ site (slow phase). Both the amplitude and the lifetime of the fast phase were affected by HL, indicating a modification of the electron transfer between Q_A_ and Q_B_, while the lifetime of the slow phase remained unchanged. In particular, the amplitudes of the slow part increased strongly in HL, again in a reversible manner (Fig. 2A). In the presence of DCMU (Fig. S2), the fluorescence decay is attributed to charge recombination between S_2_ and Q_A_^-^, with S_2_ being an oxidation state of the Mn cluster of the OEC. No difference in the fluorescence decay was seen between LL- and HL-treated cells (Fig. S2), suggesting that there is no shift in the midpoint potential of the redox couple QA/QA- and that the difference between LL- and HL-treated cells is rather due to a high reduction level of the PQ pool in HL-treated cells.

**Figure 2.**
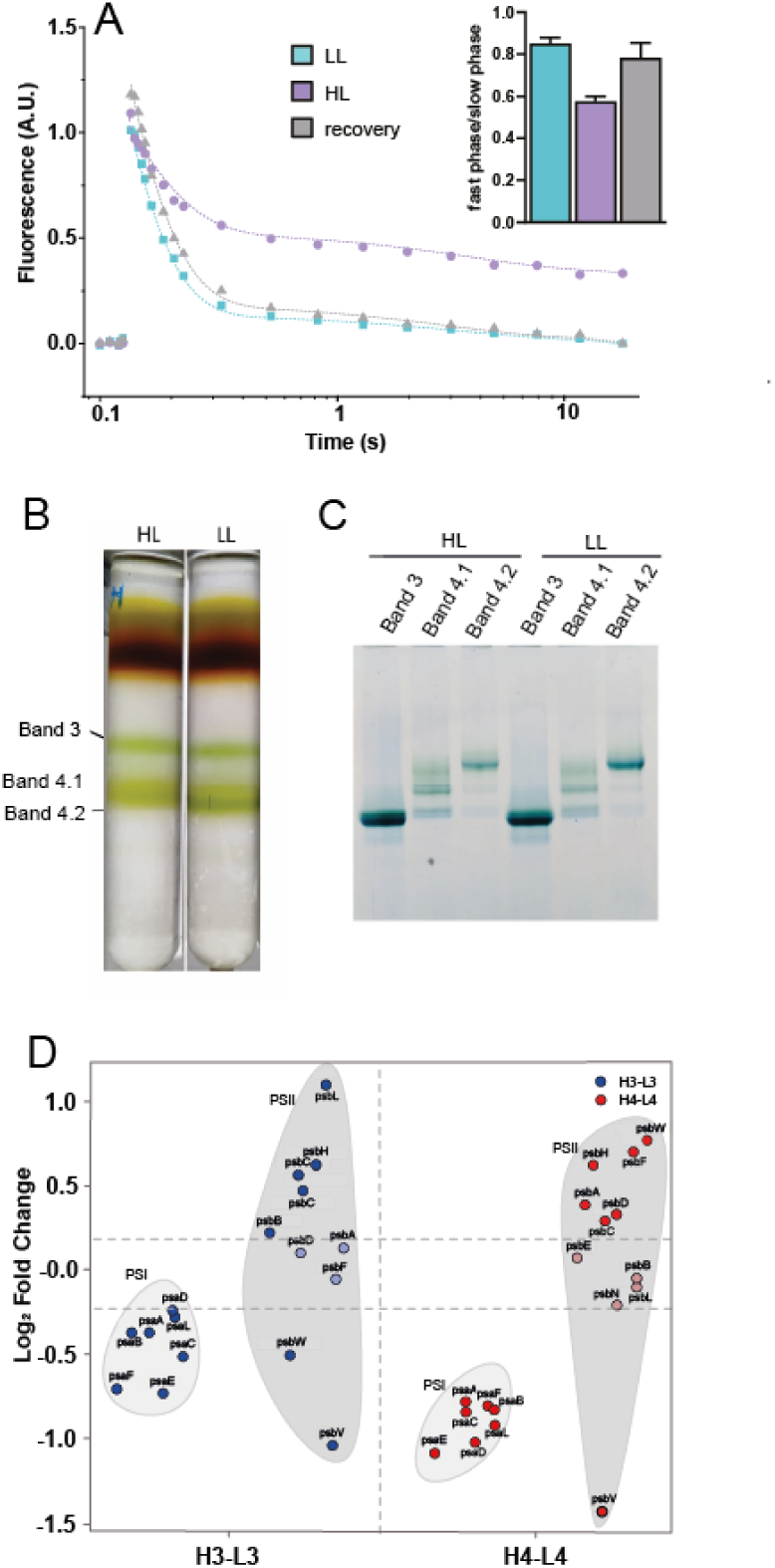
Biochemical and functional features of P. tricornutum cells exposed to different light conditions. ***A***. Fluorescence relaxation kinetics after excitation with a saturating single-turnover laser flash. The inset shows calculated amplitudes of the fast and slow phases, after fitting the traces with a double exponential decay function (dotted lines in A). **B**. Sucrose gradient of thylakoid membranes of *P. tricornutum* cells under HL and LL conditions **C**. Blue native PAGE gel of the bands from sucrose gradient (further details on Mat and Met 3). **D**. Quantitative proteomics of PSII and PSI subunits in SG bands (Log_2_FC) comparing HL and LL conditions.

Consistent with the previous observations, the effect of HL on the amplitude and the lifetime of the decay phase was rapidly reversible upon recovery to LL (Fig. S2).

To better understand the nature of the PSII changes occurring in HL, we analysed the biochemical features of LL- and HL-treated samples. Membranes were solubilized (see Methods for details) to separate the protein complexes by sucrose density gradient ultracentrifugation (Fig. 2B) and blue-native PAGE (Fig. 2C). Four major bands were observed in HL and LL (Fig. 2B) as previously described ^43^. The first band from the top to the bottom mainly contains free pigments, the second contains fucoxanthin chlorophyll a/c binding proteins (FCPs), and the last two comprise the photosystems. We further analysed bands 3 and 4 (Fig. 2B). The resolution of the sucrose gradient was sufficient to separate band 4 into bands 4.1 and 4.2. Bands 3, 4.1 and 4.2 were loaded on a blue native gel to further increase the separation and resolution of specific complexes (Fig. 2C). In-gel digestion and LC-MS/MS-based identification were used to characterize their protein content. Contrary to earlier results (showing that band 3 mainly contains PSII while PSI is mostly in band 4) ^43^, we obtained a mixture of PSII and PSI in both bands. However, differences were observed between the two light conditions: bands 3 and 4 contained higher amounts of PSI proteins under LL and higher amounts of PSII proteins under HL. Importantly, the PSII proteins that did not follow this trend: PsbV in bands 3 and 4, and PsbW in band 3, are the extrinsic subunits of the Oxygen Evolving Complex (OEC) and the PSII. These were less abundant under HL than under LL conditions (Fig. 2D), suggesting the presence of a pool of PSII monomers (Band 3) and dimers (Band 4) lacking the OEC capacity in HL. Overall, we conclude that HL treatment induces reversible structural and functional changes in PSII, with only minor alterations being observed in the overall proteome (Fig. S1).

### Topological analysis of HL-treated cells primes energy spillover between the two photosystems in diatoms

Previous work suggests that the topology of the two photosystems in the photosynthetic membranes of diatoms differ from that in vascular plants ^5^. In vascular plants, PSII is confined to the appressed grana stacks, while PSI is mainly located in the non-appressed stroma thylakoid lamellae. This physical segregation of the photosystems has a simple energetic consequence: in plants, light is absorbed and not shared directly between the two photosystems. In diatoms, no stroma and grana lamellae exist, and the two photosystems are present in the external and inner part of the three parallel membranes. This particular membrane arrangement could facilitate physical contacts between the two photosystems even in the presence of relatively small changes in the thylakoid structure, opening the way to a possible sharing of excitation energy between the two photosystems (energy spillover).

We tested this hypothesis using a spectroscopic approach ^5^. If the two photosystems are physically segregated (no spillover), blocking PSII activity with DCMU should not increase the PSI oxidation rate, while the energy absorbed by the inactive PSII should be released as chlorophyll fluorescence or thermal dissipation. Conversely, if the two photosystems are linked, blocking PSII activity with DCMU should redirect excitation energy towards PSI, leading to a higher PSI oxidation rate compared to the control. In LL-acclimated cells, no changes in the PSI oxidation kinetics were observed upon DCMU addition, ruling out that PSII fuels energy spillover to PSI (Fig. 3A, left panel). In contrast, repeating the same experiments in HL-exposed cells enhanced the PSI oxidation, as evidenced by measurements of the P_700_ oxidation rate upon DCMU addition, confirming the hypothesis of excitation energy spillover from PSII to PSI under these conditions (Fig. 3A, right panel).

**Figure 3.**
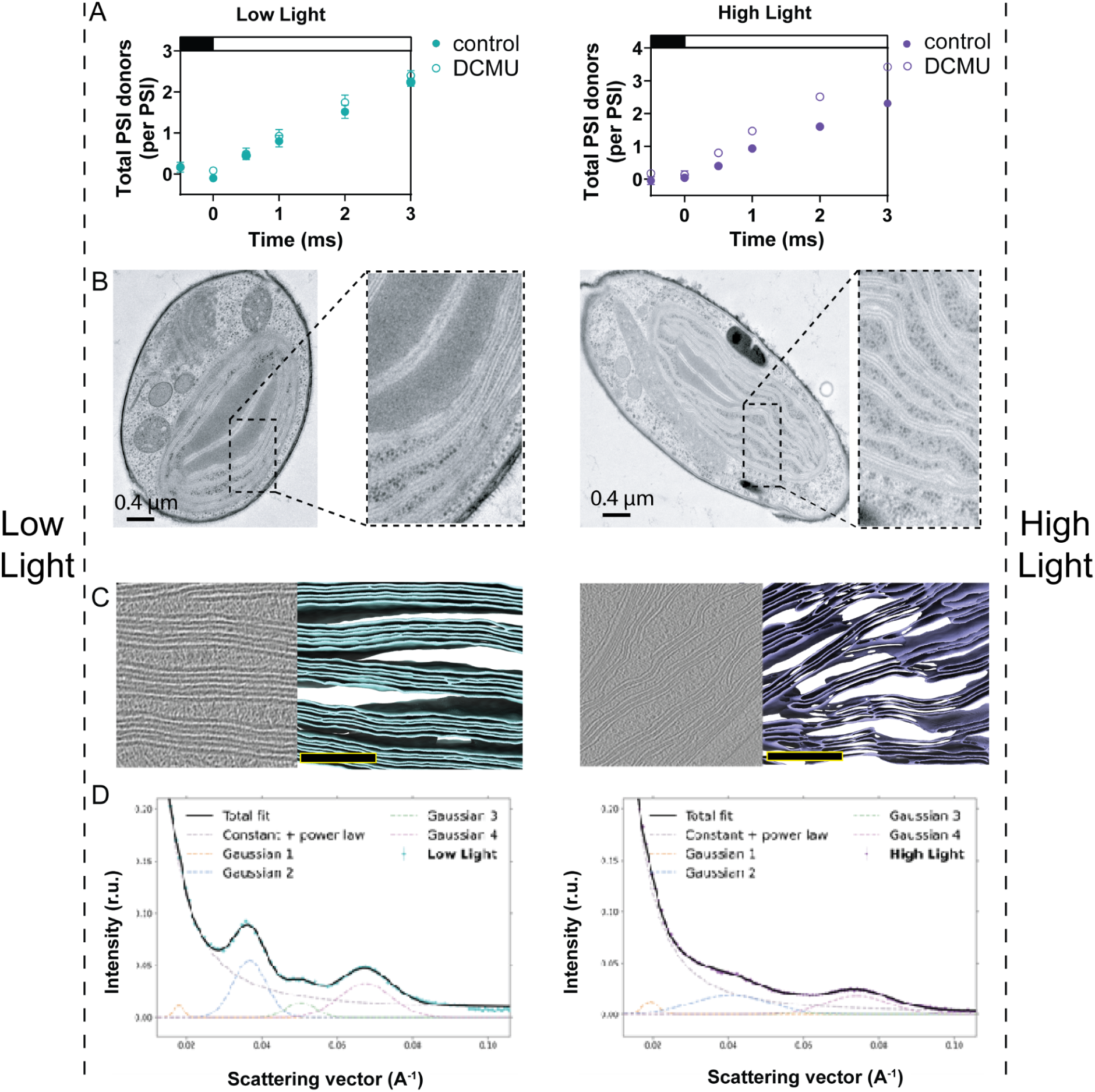
Spillover and light-induced topological changes in P. tricornutum. **A** Testing the occurrence of spillover by measuring PSI-driven electron flow *in vivo*. If the two photosystems are apart, blocking PSII activity with DCMU should not increase the PSI oxidation rate (no spillover). Conversely, if the two photosystems are linked, blocking PSII activity with DCMU should redirect excitation energy towards PSI, leading to a higher PSI oxidation rate compared to the control. **B.** Images of *P. tricornutum* cells under low light and high light obtained by transmission electron microscopy (TEM) of resin-embedded cells. **C** *In situ* cryo-electron tomography (cryo-ET) on LL- and HL-treated cells. Left images show slices through reconstructed tomograms, right colored images show 3D segmentations of thylakoid membranes (cyan: LL, purple: HL) scale bar 200 µm. **D** SANS profiles of LL (Left Panel) and HL (Right Panel) - data point of the experiment and fitted curves (continuous lines), and fitted components (dashed lines). Mathematical analysis (dotted lines) revealed the decrease of the Bragg-peak amplitude and its substantial broadening, reflecting the increased polydispersity of the membranes (Fig Similar shifts could also be recognized of the minor peaks, around 0.018 and 0.052 Å^-1^ (peak positions in LL) (Fig. 3D, see also Supplementary Table 1). Overall,the HL-induced changes in SANS point to a transition towards a more disordered thylakoid organisation^46–50^.

We consider it unlikely that changes in cyclic electron flow are not involved in the observed phenomenon. Indeed, the kinetics reported in Fig. 3A for control and DCMU poisoned samples occur in a time window (1 to 2 ms) where cyclic flow is, in principle, not operational.^44,45^

### Thylakoid membrane topology changes correlate with functional changes during the LL-to-HL transition of *P. tricornutum* cells

As light acclimation modifies the cellular features of diatoms ^6^, with apparent reorganization of the photosynthetic complexes within the membranes, we investigated the structural consequences of light regime on plastidial thylakoids, looking for possible modifications that could predispose the system to spillover.

Both conventional transmission electron microscopy (TEM; Fig. 3B) and *in situ* cryo-electron tomography (cryo-ET; Fig. 3C) data support this hypothesis. TEM of resin-embedded sections showed that general morphological features of the chloroplast were not substantially altered by the treatment, suggesting that short-term HL exposure does not cause substantial dismantling of the thylakoid network. However, both EM and cryo-ET revealed less ordered thylakoid membranes upon HL transition: stacks of three parallel thylakoids were most commonly observed under LL conditions, whereas more irregular stacks of undulating membranes appeared under HL, suggesting that this treatment affects the membrane ultrastructure (see also Fig. S3 and S4 and supplementary videos 1-4).

We confirmed these findings via Small-Angle Neutron Scattering (SANS) analysis (Fig. 3D), which provides a readout of how the bulk membrane organization changes *in vivo* within a suspension of many cells. SANS is a non-invasive experimental tool which provides structural information averaged over the entire sample volume. This analysis showed well discernible modifications in the two characteristic peaks reflecting the thylakoid repeat distance and periodic order. LL cells (Fig. 3D, Left Panel), in good accordance with earlier SANS profiles recorded under similar conditions, displayed two major peaks at around 0.037 and 0.065 Å^-1^; the former, so-called Bragg peak, has been shown to originate from the thylakoid repeat distance of about 170 Å; whereas the second, major high-Q peak was proposed to arise from adjacent membrane pairs (Nagy et al. 2012)^46–50^. Upon exposing the cells to HL for 2 hours, the Bragg peak almost vanished (Fig. 3D, Right Panel), consistent with a substantial weakening of the periodic order of the thylakoids. The ability to detect the same membrane reorganization with SANS not only confirms our TEM and cryo-ET observations, but also indicates that these ultrastructural changes occur widely within the population of cells.

Finally, a time course analysis of spillover recovery (Fig. S5A) and RC-quenching recovery (Fig. S5B) after 2h of HL exposure suggests that, upon induction, the two processes can follow similar relaxation kinetics. This finding strongly suggests their concomitant presence in living cells.

## Discussion

In this study, we have revisited the molecular bases of photoprotection in diatoms. To this end, we developed an easy experimental protocol to test the role of different photoprotective responses in *P. tricornutum*. Cells were grown in LL and then exposed to a sudden increase in light intensity. The duration of the treatment was short enough to not induce constitutive adaptive changes to light of the photosynthetic apparatus ^6^, but the extent of the light change was strong enough (∼9-fold increase) to trigger short-term photoprotective responses.

We observed that some photoprotective responses display characteristics different from the ‘classic’ NPQ process. Despite the presence of classic NPQ signature features (carotenoid de-epoxidation) in our samples, some biochemical and functional features (the simultaneous decrease in NPQ, Fv/Fm and oxygen evolution) were consistent with a transient, reversible photoinhibition (via charge recombination). The partial loss of Fv/Fm and the concomitant partial PSII disassembly, possibly to monomeric form, are compatible with so-called ‘reversible’ photoinhibition, a condition in which PSII can no longer contribute to forward electron flow, but it is still able to rapidly recover activity in the dark without protein biosynthesis ^39–41^. At the same time, we observed changes in the thylakoid membrane topology that could promote proximity between the two photosystems in HL, when the parallel arrangement of the thylakoids is perturbed. In particular, we pinpointed regions where the two photosystems could be very close and therefore could physically interact. This is the prerequisite to obtain energy spillover from PSII to PSI. This phenomenon can also be considered an alternative form of PSII energy quenching by the nearby PSI reaction centres. Overall, these two PSII-localized energy quenching mechanisms could coexist with qE, given their comparable kinetics of onset and relaxation (Fig. 4).

**Figure 4.**
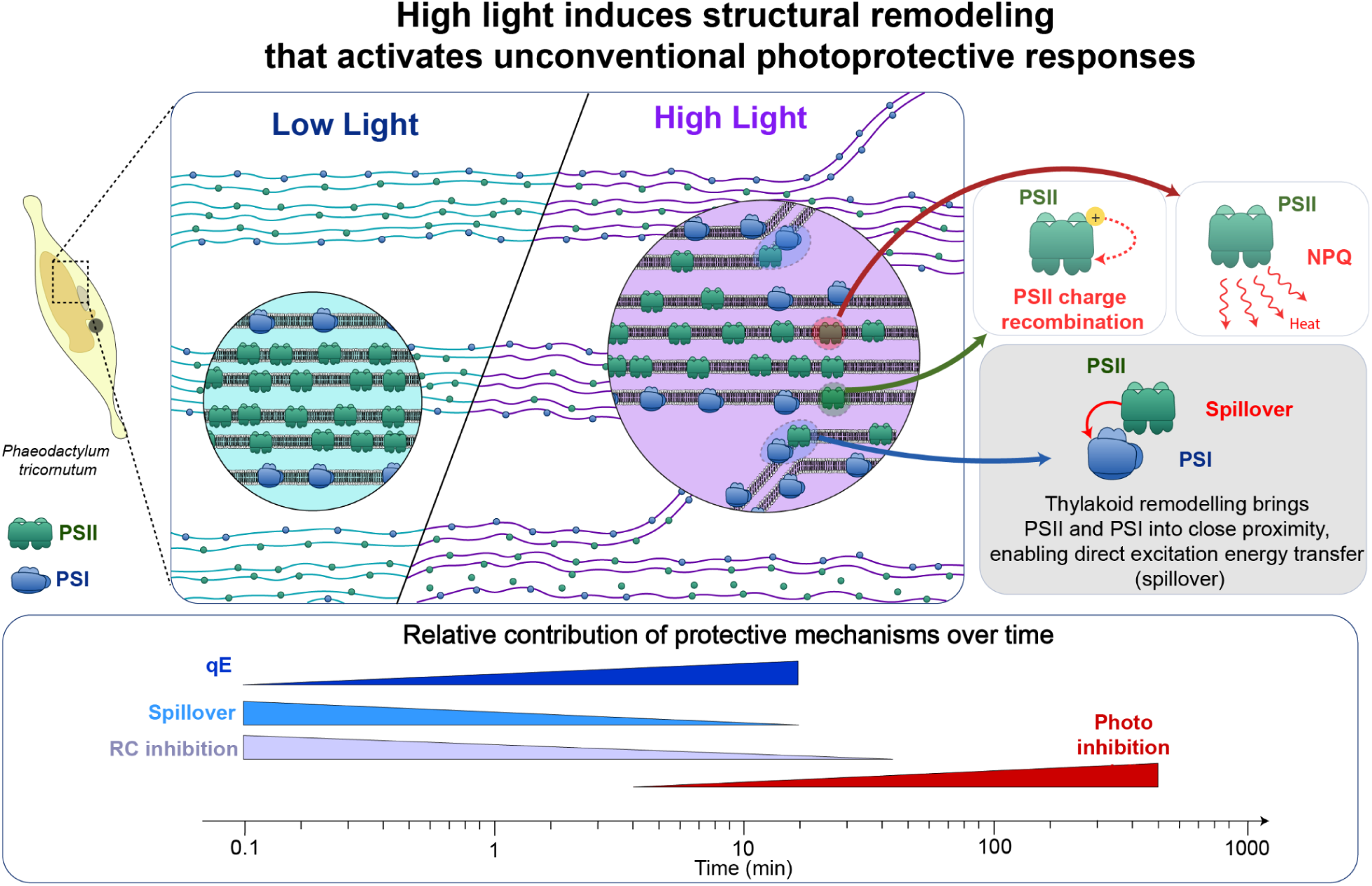
High-light-induced thylakoid structural remodelling and associated photoprotective responses in Phaeodactylum tricornutum. Top left: thylakoid organization under low light (ordered stacks, with PSII in green and PSI in blue) and high light (remodelled, less ordered membranes allowing PSII-PSI contacts). Top right: photoprotective processes associated with this reorganization, namely excitation energy spillover from PSII to PSI, accumulation of the quencher P680^+^ in the reaction center and charge recombination in PSII and non-photochemical quenching (NPQ). Bottom: conceptual representation (not to scale) of the relative contribution of energy dependent quenching (qE), spillover, reaction-centre (RC) inhibition and photoinhibition as a function of time after the onset of high light.

Beyond qE, which is an extensively described process ^51^, all the dissipation processes identified in this work have already been observed in other systems. Energy dissipation via charge recombination RC-NPQ has been characterized by spectroscopic analysis in purified systems ^52–54^, not only as a side effect of the process of charge separation but also as a possible way to lower the photochemical efficiency. In diatoms, its occurrence has been documented and likely reflects P_680_^+^Q_A_^-^ charge recombination within PSII^18^.

Finally, the existence of energy spillover between the two photosystems is documented in several phototrophs. A faster PSI turnover was measured upon Q_A_ reduction in the red microalga *Porphyridium cruentum* ^55–56^, whose unstacked phycobilisome-decorated thylakoids do not show any apparent segregation between PSI- and PSII-rich domains. A faster PSI turnover was also shown in a red macroalga, *Chondrus crispus,* under similar conditions, resulting in an increased capacity to oxidise plastocyanin (the direct electron donor to PSI) ^50^. In general, spillover is restricted to particular conditions in plants^57^, because the two photosystems are physically segregated between two compartments: the grana stack and the stroma lamellae. However, Bag et al. (2020) ^58^ combined ultrafast fluorescence spectroscopy and EM in conifers to show thylakoids structural degeneration at low temperatures. This structural rearrangement promotes constitutive PSII-PSI contact and sustained quenching, providing photoprotection against early spring sunlight after severe cold winters ^58^. Red macroalgae, in which no strong PSII-PSI physical segregation is observed, can also develop sustained spillover to protect PSII under high light ^55,59^. Finally, energy spillover in response to environmental stress has been proposed in the symbiotic alga *Symbiodinium* ^60^.

Altogether, the examples above demonstrate that excitation energy spillover between PSII and PSI occurs across a broad phylogenetic range of phototrophs. Yet, in most cases this process is either constitutive (as in organisms lacking strong lateral heterogeneity of their thylakoid membranes) or induced by long term stress conditions (as in overwintering conifers)^61^. The situation we describe in *P. tricornutum* is peculiar in several important respects. First, spillover is rapidly induced by a moderate light shift (∼9-fold), and rapidly reversed under conditions that do not cause irreversible photodamage. Second, it operates in parallel with at least two other short-term photoprotective dissipative mechanisms, namely RC-NPQ and canonical qE, providing a layered and tuneable global photoprotective response. Third, the structural basis for spillover in diatoms probably resides in the reversible remodelling of thylakoid membrane topology, which opens the way to a less strict lateral heterogeneity of the complexes. Overall, the reversibility, rapidity, and dynamics of these quenching processes may constitute a key adaptive advantage in their rapidly fluctuating light environments.

Although this phenomenon occurs in vascular plants under specific conditions, energy spillover appears to systematically occur in phototrophs with less strict lateral heterogeneity of thylakoid membrane organization favouring physical proximity between the two photosystems, as is the case in marine plankton. Since the PSII to PSI stoichiometry in diatoms is ≥1 ^62,63^, the monomerization of a fraction of PSII complexes as observed here would not limit linear electron flow, which requires an equimolar ratio between the two photosystems. Moreover, the presence of excess PSII and the possibility of transient interaction between the two photosystems would ensure flexible modulation of light acclimation responses, combining excitation energy spillover and PSII reaction centre quenching. Both are relatively fast processes, possibly with different rise times, that do not require gene expression or de novo protein synthesis, but only a modification of inter-photosystem contacts or the selective disassembly of specific PSII subunits. It is therefore tempting to propose that both processes represent physiological photoprotective responses in other oceanic plankton members, since the higher flexibility of the thylakoid membrane’s lateral segregation seems to be conserved. Their further characterization should provide a more comprehensive understanding of the dynamics of photosynthetic regulation in the ocean (Fig. 4).

To conclude, we demonstrate that multiple strategies to deactivate light excitation pressure on PSII can occur simultaneously in living diatoms. They act on a similar time scale suggesting the existence of a cause-effect relationship.

## Materials and methods

### 1 Cell culture and growth conditions

The *P. tricornutum* Pt1 strain (CCAP 1055/3) was obtained from the Culture Collection of Algae and Protozoa, Scottish Marine Institute, UK. Cells were grown in the ESAW (Enriched Seawater Artificial Water) medium ^64^ supplemented with NaNO_3_ (468 mg l^-1^) and NaH_2_PO_4_·H_2_O (30 mg l^-1^), in 50 mL flasks in a growth cabinet (Infors HT Incubator, Bottmingen, Switzerland), at 19 °C, with light intensity of 40 µmol photons m^-2^ s^-1^, and a 12-h light/12-h dark photoperiod. High light treatments were performed by irradiating the cells with 350 µmol photons m^-2^ s^-1^ for 2 hours without rotation after the onset of light, with the same light sources. Temperature was maintained at 19 °C. The recovery treatment consisted of placing cells that had undergone high light treatment in low light conditions immediately afterwards for a period of 2 hours without rotation and at 19 °C.

### 2 Thylakoid solubilization and sucrose gradient fractionation

1 L of *P. tricornutum* cell culture at a concentration of 5 × 10⁶ cells mL⁻¹ was concentrated to 5 × 10⁸ cells mL⁻¹ using 50 mL tubes and a centrifuge at 600 × g for 3 min at 19 °C. The concentration was adjusted using a 0.4 M sorbitol buffer (it is important to maintain a temperature close to the culture temperature until cells disruption). The sample was then placed in 2 mL tubes with secure caps (500 µL per tube) containing 500 mg of glass beads (425-600 µm, Sigma Aldrich, G8772) and then placed in a cell destructor (Precellys Evolution Touch, Bertin Technologies) in a cold chamber (4 °C) at 10,000 rpm for three cycles of 30 seconds of agitation and 30 seconds of pause (sample must be at 4 °C and in darkness for the rest of the experiment). The sample was recovered (beads should not be recovered; the use of a small micropipette tip is recommended) and then centrifuged at 1000 × g for 1 min at 4 °C. The supernatant and pellet were separated, the pellet was mixed with 500 µL of 0 M buffer and the two solutions were centrifuged at 3500 × g for 10 min at 4 °C. The supernatants were discarded and the pellets were combined with 800 µL of 0.1M sucrose buffer and 1% final αDDM (0M and 0.1M buffers contained 0M sucrose and 0.1M sucrose respectively with addition of some salts (5mM MgCl2, 22mM KCl, 10mM NaCl and 20mM HEPES at pH8 (KOH)). The sample was gently rotated for 10 minutes at 4 °C, then centrifuged at 20,000g for 20 minutes, still at 4 °C. The supernatant was collected and placed on the sucrose gradient.

For the sucrose gradient, we used high-speed centrifugation tubes (Ultra-Clear, 14 × 89mm, 344059, Beckman Coulter), which were first filled halfway with a 10% sucrose solution and then halfway with a 40% sucrose solution. The tubes were capped and placed on a homogenization rack (Gradient Master 108, BioComp) to create a 10-40% linear sucrose gradient. 400 µL of sample were loaded on the sucrose gradient and centrifuged at 39,000 rpm, 4 °C, and 20 µmHg vacuum for 18 hours using an ultracentrifuge (Optima L-90K Ultracentrifuge, Beckman Coulter) and a support (SW 41 Ti Swinging-Bucket Rotor, Beckman Coulter). After centrifugation, the bands were recovered on a 96-well plate using a fractionator (Piston Gradient Fractionator, BioComp): speed 0.2 mm/s, distance 1 mm corresponding to approximately 150µL fractions.

### 3 Blue Native PAGE and in-gel digestion

The bands recovered from the 96-well plate after sucrose gradient separation were pooled and concentrated using 100 kDa filters. Each of the bands was introduced into a precast Blue Native PAGE gel (Native PAGE 3-12% Bis-Tris, 1.0mm, 10-well, BN1001BOX, Invitrogen) and migrated according to the protocol ^65^. Alternatively, two 150µL SG bands were used for in solution digestion as previously ^66^. Briefly, samples were reduced (DTT 10 mM at 37°C for 30 min), alkylated (20 mM iodoacetamide RT in the dark for 1 hour) and digested (trypsin 1:20 ratio overnight at 37 °C, T1426, Merck). TFA 1% final concentration was added to stop digestion and centrifugation at 21,000 × g to get rid of sodium deoxycholate. A desalting step was performed on Oasis®Prime HLB µElution plate (96-wells, Waters Corporation, 186008052). Samples were acidified with TFA (pH 2), then columns were washed twice with 200 µL of pure ACN and twice with 200 µL of 0.1% TFA. Then the sample was passed through the column and was washed twice with 200 µL of 0.1% TFA before final elution with twice 35 µL of a solution containing 50% ACN and 0.1% TFA. Samples were then acidified (pH 2), dried in a vacuum concentrator and stored at -20 °C.

### 4 LC-MS/MS sample preparation and analysis

For quantitative proteomics on whole cells, we collected ∼1 × 10^7^ cells by centrifugation at 1000 × g for 3 min right after the light treatment, lysed on a Precellys and precipitated by adding ice-cold pure acetone in steps of 50%, 50%, 100%, 100% and 200% of the initial solution, under constant stirring at 600 rpm, at RT and for 5 min between additions. Samples were stored at -20 °C overnight and centrifuged at 21,000 × g for 30 min at 4 °C and the acetone was discarded by pouring and then drying under a fume hood at RT for at least 15 min. For the comparison of the sucrose gradient fractions bands were harvested using BioComp Piston Gradient Fractionator (PGF)(BioComp, Canada).

Pellet was then resuspended with a solution with final 1% w/v sodium deoxycholate (SDC) and 0.1 M Tris (pH 9) using mechanical assistance (2 glass beads of 4 mm in a 2 mL tube at 1000 rpm for 2 hours). Protein concentration was estimated using Bradford protein assay (Protein Assay Dye Reagent Concentrate, BioRad, 5000002) following the manufacturer’s recommendations. Samples were reduced (DTT 10 mM at 37°C for 30 min), alkylated (20 mM iodoacetamide RT in the dark for 1 hour) and digested (trypsin 1:20 ratio overnight at 37 °C, T1426, Merck). TFA 1% final concentration was added to stop digestion and centrifugation at 21,000 × g to get rid of sodium deoxycholate. A desalting step was performed on Oasis®HLB 96-well Plate 30 µm (Waters Corporation, WAT058951). Samples were acidified with TFA (pH 2), then columns were washed twice with 1 mL of pure ACN and twice with 1 mL of 0.1% TFA. Then the sample was passed through the column and was washed twice with 1 mL of 0.1% TFA before final elution with twice 300 µL of a solution containing 80% ACN and 0.1% TFA. Samples were then acidified (pH 2), dried in a speed vacuum and stored at -20 °C.

Samples were resuspended in 5 % acetonitrile and 0.1 % TFA and analyzed using nano-liquid chromatography coupled with tandem mass spectrometry (nano-LC-MS/MS). MS and MS/MS data were acquired in data-dependent acquisition mode using a Dionex Ultimate 3000 RSLCnano and Q-Exactive HF respectively (both Thermo Fisher Scientific). Peptides were sampled on a 300 μm × 5 mm PepMap C18 precolumn (Thermo Fisher Scientific) and separated on a 75 µm × 250 mm Aurora Ultimate C18 column (IonOpticks). The nano-LC utilized a 140-minute gradient (2–41 % acetonitrile in 0.1 % formic acid) at a flow rate of 300 nL/min. Full MS spectra (m/z 400–1600) were acquired at a resolution of 60,000, and MS/MS spectra were acquired at a resolution of 15,000 using a stepped normalized collision energy of 30 eV.

Raw data were processed using FragPipe 22.0 ^67^ for peptide identification and using UniProt database ^68^ (Phaeodactylum tricornutum taxonomy, downloaded 25 January 2025), a classical contaminants database, and their corresponding reversed databases. Trypsin was chosen as the cleavage enzyme, with a maximum of three missed cleavages allowed. The allowed peptide modifications were: (1) acetylation of the protein’s N-terminus (variable); (2) methionine oxidation (variable) and (3) cysteine carbamidomethylation (fixed). Precursor and fragment mass error tolerances were set to 20 ppm. Data were statistically analysed using ProStaR (version 1.38.1) ^69^. Proteins identified as contaminants were discarded. Data were log_2_ transformed and statistical significance between high light and low light samples was verified using the limma test ^70^. Differentially abundant proteins were selected using a p-value cut-off providing an FDR of less than 5% according to the Benjamini–Hochberg procedure ^71^.

### 5 Fluorescence-based measurements

Light-induced fluorescence kinetics were measured using a fluorescence CCD camera recorder (Speedzen imaging system, BeamBio, France) as described ^72^ on cells in exponential phase, at a density of 2–3 × 10^6^ cells mL^-1^. Photochemical activity and structural dynamics of PSII reaction centres (Fv/Fm) and NPQ were calculated, respectively, as (Fm-F0)/Fm and (Fm-Fm’)/Fm’ ^73^ where Fm and Fm’ are the maximum fluorescence emission levels in the dark and light-acclimated cells, measured with a saturating pulse of light. All samples were adapted to dim light (<10 µmol photons m^-2^ s^-1^) for 15 min before measurements. The maximal NPQ response was measured upon exposure for 7.5 min to saturating red light of 600 µmol photons m^-2^ s^-1^. Measurements shown in Fig. 1A and B are representative of a total of 3 biological samples. For the experiments in Fig. 2A, cells were exposed to single turnover saturating flashes (duration ∼7 nanoseconds), provided by a laser dye (L2S698) pumped by a frequency doubled Nd-YAG source (Quantel, France) and chlorophyll fluorescence has been recorded with a JTS-10 (Biologic, France).

### 6 Spectroscopic measurements

Energy spillover from PSII to PSI, redox changes of P_700_ and of its electron donor pool were monitored as in Flori et al. 2017 ^5^, using a JTS-10 spectrophotometer (Biologic, France). Cells concentrated at 2 × 10^7^ cells mL^-1^ were placed under two light conditions: high light (350 µmol photons m^-2^ s^-1^) and low light (40 µmol photons m^-2^ s^-1^) for 2 hours without rotation after the onset of light at 19 °C. Measurements were taken in cuvettes containing 1.5 mL of P. tricornutum solution with a light intensity of 590 µmol photons m^-2^ s^-1^ at 705 nm. After the first measurement, the first PSII electron transport inhibitor DCMU (3-(3,4-dichlorophenyl)-1,1-dimethylurea) was added to the cuvette at a final concentration of 20 µM, before performing a second measurement on the JTS-10 with the same light parameters. A third measurement with the same light parameters was finally performed after adding a second photosynthesis inhibitor (hydroxylamine at a final concentration of 2 mM). Five repetitions were performed on two independent biological replicates.

### 7 Pigment analysis by high-performance liquid chromatography (HPLC)

Pigment analysis was performed as described in Allorent, G. et al ^74^ from cells harvested in mid-exponential phase.

### 8 Oxygen measurements

Oxygen measurements were taken on 700 µL of P. tricornutum culture concentrated at 1.5 × 10^7^ cells/mL using a Photosynthesis Yield Analyzer (Mini-PAM II, Walz) connected to a Retractable Fiber Oxygen Microsensor (OXR50-OI, Pyroscience) and a stirrer (Mini MR standard, IKA). Oxygen and light data were processed using WinControl3 software. The light protocol was as follows: (1) darkness followed by a measuring light and a saturating pulse (when measuring light is stable) before turning off the measuring light; (2) dim actinic light (48 µmol photons m^-2^ s^-1^) followed by a measuring light and a saturating pulse (when measuring light is stable) before turning off the measuring light; (3) moderate actinic light (298 µmol photons m^-2^ s^-1^) followed by a measuring light and a saturating pulse (when measuring light is stable) before turning off the measuring light; (4) strong actinic light (1200 µmol photons m^-2^ s^-1^) followed by a measuring light and a saturating pulse (when measuring light is stable) before turning off the measuring light. Only dim and moderate light has been used for data interpretation as the strong actinic light of 1200 µmol photons m^-2^ s^-1^ lies outside the range discussed in this article.

### 9 Transmission electron microscopy

Cells of *P. tricornutum* were fixed in a double-strength fixative (4% (w/v) formaldehyde, 0.4% (v/v) glutaraldehyde) in PHEM buffer (PIPES 60 mM, HEPES 25 mM, EGTA 10 mM, MgCl_2_ 2 mM; pH 7.0) in an equal volume to the culture medium (ESAW), and then diluted into a standard strength fixative (2% (w/v) formaldehyde (EMS, USA) and 0.2% (v/v) glutaraldehyde (EMS, USA)). After 15 min, fresh standard strength fixative was replaced and fixation proceeded for 30 min at 20 °C, under agitation. Cells were washed three times with 50mM glycine in PHEM buffer and after centrifugation were embedded in 12% gelatin in PHEM. The gelatin-embedded blocks were cryoprotected in 2.3 M sucrose in rotating vials at 4 °C (16 h). Samples vitrification was obtained in liquid nitrogen following the plunge and freezing technique ^65^. Thin sections (80 nm) were prepared at -110 °C with a diamond knife (Diatome, Switzerland). Ribbons were picked-up with a drop of 1% (w/v) methylcellulose/1.15 M sucrose in PHEM buffer. Sections were thawed and transferred to Formvar carbon-coated nickel grids.

### 10 Small-angle Neutron Scattering (SANS)

Experiments were performed on the D22 SANS instrument at the ILL (Institut Laue-Langevin) in Grenoble, France, where a high-flux research reactor provides a continuous neutron beam (http://www.ill.eu/instruments-support/instruments-groups/instruments/d22/). Sample suspensions filled in 2-mm quartz cuvettes were mounted in a sample holder placed in the neutron beam, defined by a 10 mm × 6 mm size aperture in front of the sample. Neutrons scattered from the sample were detected with a position-sensitive detector. Data acquisition was performed as described previously ^47,75^. The 1D scattering curve was fitted with a linear combination of a constant, a power-law term, and four Gaussian functions in the q-range of 0.012-0.106 Å⁻¹. [GN1] The thylakoid membrane repeat distances (RDs) were determined from the center positions (q_0,2_) of the second peak (found between about 0.037 and 0.043 Å⁻¹) as RD = 2π/q_0,2_.Fitting results for all the parameters in Eq(1) for the SANS curves on the different low and high light grown samples. where *I(q)* is the scattering intensity, *q* is the scattering vector, *i_0_* is the constant background, *Aq^p^* is the power-law background, *A_k_*the amplitude of the *k*-th Gaussian peak, *q_0,k_* is the position of the *k*-th Gaussian peak, σ_k_ is the standard deviation of the *k*-th Gaussian peak. (For clarity, the errors of the fitted parameters, which typically were smaller than 5%, are not displayed.)

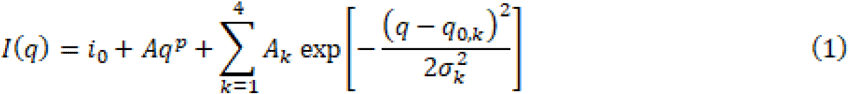

### 11 Cryo-ET sample preparation and data acquisition

*P. tricornutum* was grown to an exponential phase in low-light conditions (40 µmol photons m^-2^ s^-1^) prior to vitrification. Cells were concentrated at 800 × g for 5 min and 4 µL of cell suspension was applied on 200-mesh R ¼ SiO_2_-film covered gold grids (Quantifoil Micro Tools) and plunge frozen using a Vitrobot Mark IV (Thermo Fisher Scientific). EM grids were clipped into Autogrid supports (Thermo Fischer Scientific) and loaded into Aquilos 1 or 2 FIB-SEM instruments (Thermo Fischer Scientific), where they were thinned with a Gallium ion beam. The resulting EM grids with thin lamellae were transferred to a transmission electron microscope for tomographic imaging. Cryo-ET data were acquired on a 300 kV Titan Krios G2 microscope (Thermo Fisher Scientific), equipped with a post-column energy filter (Quantum, Gatan) and a direct detector camera (K2 summit, Gatan), as well as on a 300 kV Titan Krios G3i microscope (Thermo Fisher Scientific) equipped with a BioQuantum post-column energy filter (Gatan) and a K3 direct electron detector (Gatan). Each image was recorded in counting mode with ten frames per second. The target defocus of individual tilt-series ranged from −2 to −5 µm. Total dose per tilt series was approximately 120 e⁻/Å² . Data were acquired in EER mode with a calibrated image pixel size of 2.93 Å.

### 12 Cryo-ET data analysis

TOMOMAN MATLAB scripts (https://github.com/williamnwan/TOMOMAN/)^67^ (version 0.6.9) were used to pre-process the tomographic tilt series data. Raw frames were aligned using MotionCor2 (version 1.5.0) ^76^, then tilt-series were dose-weighted ^77^ followed by manual removal of bad tilts. The resulting tilt-series (binned 4 times) were aligned in IMOD (version 4.11) ^78^ using patch tracking and were reconstructed by weighted back projection. Cryo-CARE (version 0.2.1)^79^ was applied on reconstructed tomogram pairs from odd and even raw frames to enhance contrast and remove noise. Snapshots of denoised tomograms were captured using the IMOD 3dmod viewer. Denoised tomograms were used as input for automatic segmentation using MemBrain-Seg^80^. The resulting segmentations were manually curated in Amira (version 2021.2).

## Acknowledgements.

The authors thank the ERC ChloroMito (833184) for funding. We thank Stéphane Tachon and Anna Sartori-Rupp from the NanoImaging Core Facility, Institut Pasteur, 25-28 Rue du Dr Roux, 75015 Paris, France for discussions and help with cryo-ET sample preparation.

S.F. acknowledges support from EMBO short-term fellowship (8015), a Royal Society international exchange grant (IESR3183037) and an EIPOD-LinC postdoctoral fellowship from EMBL. G.N. acknowledges resources from the Spallation Neutron Source, a DOE Office of Science User Facility operated by the Oak Ridge National Laboratory. The authors acknowledge the Institut Laue-Langevin (ILL, Grenoble, France) for the allocation of beamtime on the D22 SANS instrument and thank Lionel Porcar for assistance with the instrument setup. The work of R.U. was financed from the OTKA PD 138540 grant. G.G. received support from a grant from the European Union under the LERCO project (CZ.10.03.01/00/22_003/0000003) via the Operational Programme Just Transition.

## Author contributions

All authors contributed to this manuscript and approved the final submitted version. G.F., P.A. and S.F. conceptualized and directed the project. D.P.C., D.T., C.U., C.G., L.P., D.F., C.M., W.W., G.G., R.U., G.N., P.J., G.S., A.K.-L, B.D.E., P.A., G.F. and S.F. carried out the experiments. D.P.C., G.F., P.A., R.U. and S.F. analyzed the data. D.P.C., B.D.E., G.F., P.A. and S.F. wrote parts of the draft and contributed to critical revision of the paper.

## Funding

We acknowledge funding from the European Research Council (ERC) for the Chloro-Mito project (grant no. 833184 to G.F., S.F.) and for the PlanktON project (grant no. 101099192 to G.F. and P.A.) from the European Innovation Council (EIC). P.A. also acknowledges support from the European Union’s Horizon 2020 research and innovation programme via the Marie Skłodowska-Curie grant agreement no. 101066400 (PhotoLINK) and the 2022 ANR PRC program (PHOTO_DYN—ANR-22-CE20-0015-01). Proteomic experiments were partially supported by Agence Nationale de la Recherche under projects ProFI (Proteomics French Infrastructure, ANR-10-INBS-08; ANR-24-INBS-0015). This work was also supported by GRAL ANR-17-EURE-0003 and/or ANR-10-LABX-49-01.

## Notes

### Competing Interest Statement

The authors have declared no competing interest.

