## Supplementary figures for "Thylakoid plasticity primes photoprotection in diatoms"

### Supplementary material

**Figure S1. Net photosynthesis rate at different light intensity**

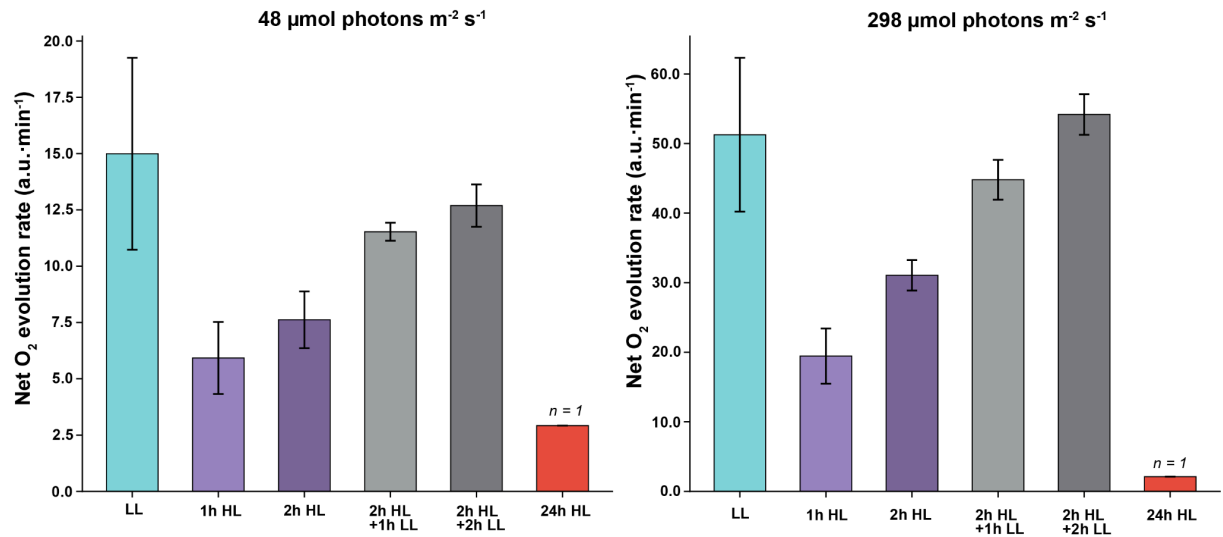

**Figure 1D. Net photosynthesis rates measured at two actinic light intensities.** Oxygen evolution was measured in *P. tricornutum* cells using a MiniPAM II under low (48 μmol photons m<sup>-2</sup> s<sup>-1</sup>, left panel) and high (298 μmol photons m<sup>-2</sup> s<sup>-1</sup>, right panel) actinic light. Rates are expressed as the slope of the oxygen trace during the linear phase of O<sub>2</sub> evolution at each light intensity (arbitrary units per minute). Cells were either maintained under low light (LL, 40 μmol photons m<sup>-2</sup> s<sup>-1</sup>), shifted to high light (HL, 350 μmol photons m<sup>-2</sup> s<sup>-1</sup>) for 1 or 2 hours, allowed to recover in LL for 1 or 2 hours after a 2 h HL exposure, or maintained under HL for 24 hours. HL exposure led to a decrease in net O<sub>2</sub> evolution at both measuring intensities, which was reversible upon return to LL. Prolonged HL exposure (24 h) virtually abolished photosynthetic activity. Error bars represent the standard deviation from two independent experiments (n = 2); the 24 h HL condition was measured in a single experiment (n = 1). Data can be found in Supp. Data. Net photosynthesis

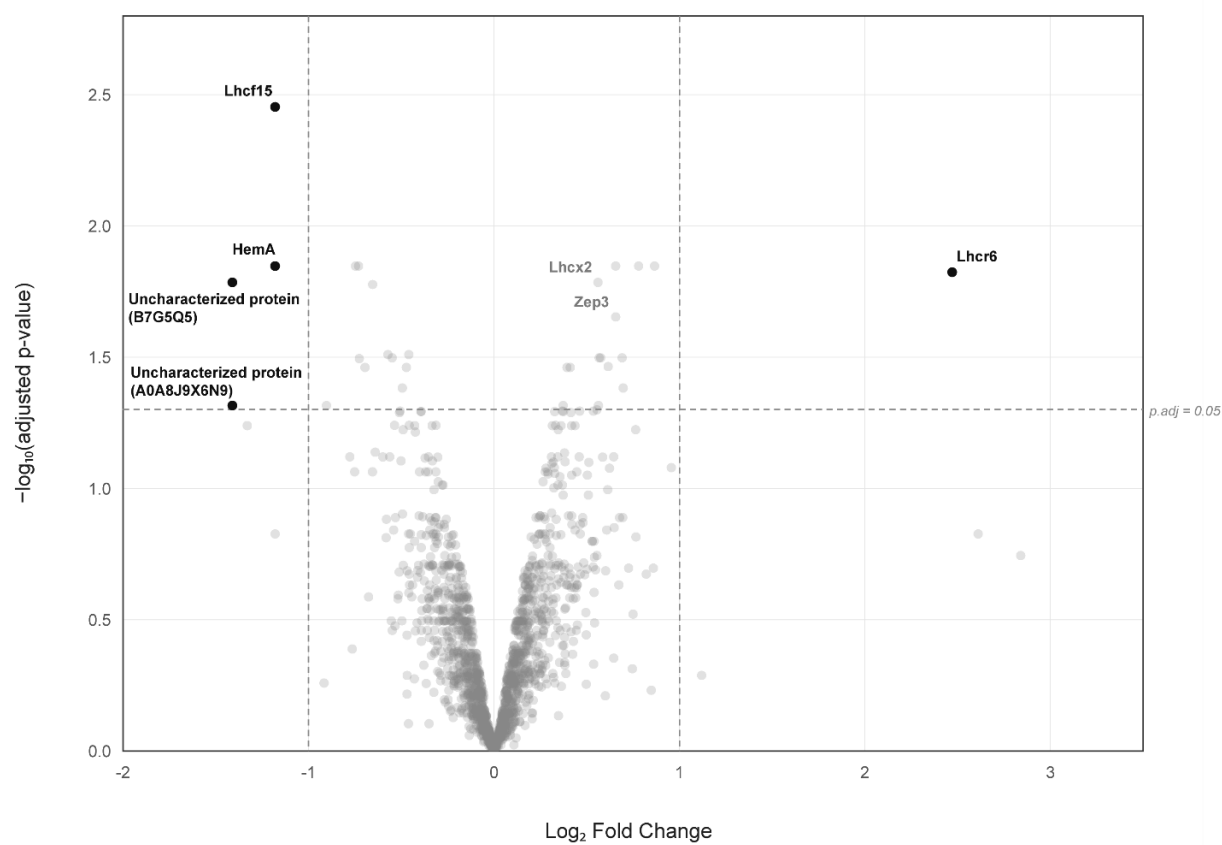

**Figure S2. Differential protein accumulation upon 2h of high light treatment.**

Volcano plot of protein abundance changes in whole-cell lysates after 2 h of HL versus LL ($n = 3$, data represent the average intensities of replicates). $\text{Log}_2\text{FC}$ calculated using the intensity values between HL versus LL and adjusted p-values calculated using the Limma test in ProStaR. Significance thresholds were set at a $\text{Log}_2\text{FC}$ over 1 and an adjusted p-value of 0.05 (dashed lines). Proteins passing both thresholds are labelled in black; light-acclimation proteins significant at adjusted $p < 0.05$ but below the fold-change threshold (Lhcx2, Zep3) are labelled in grey.

A. Low-Light

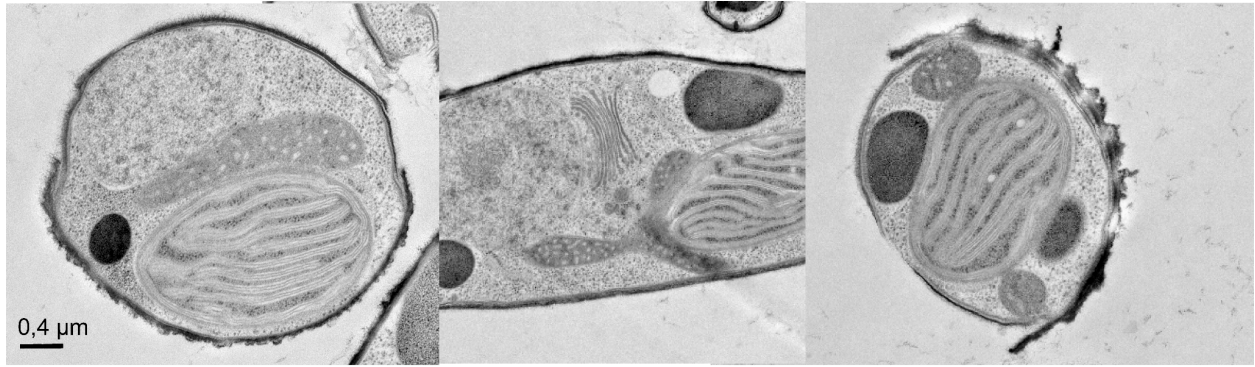

B. High-Light

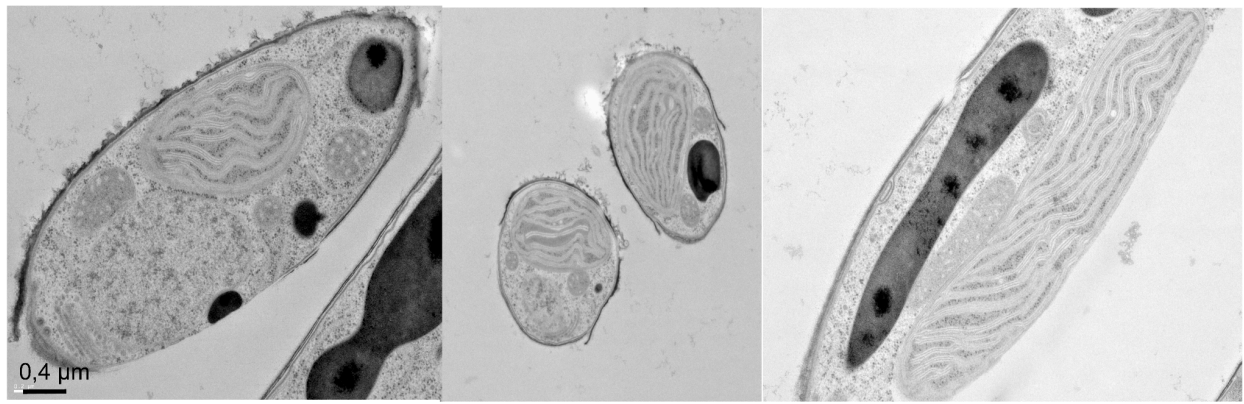

**Figure S3. Gallery of TEM images of LL (left) and HL (right) *P. tricornutum* cells.** Images of *P. tricornutum* cells under low light and high light obtained by transmission electron microscopy (TEM) of resin-embedded cells. Same conditions as in Fig. 3B.

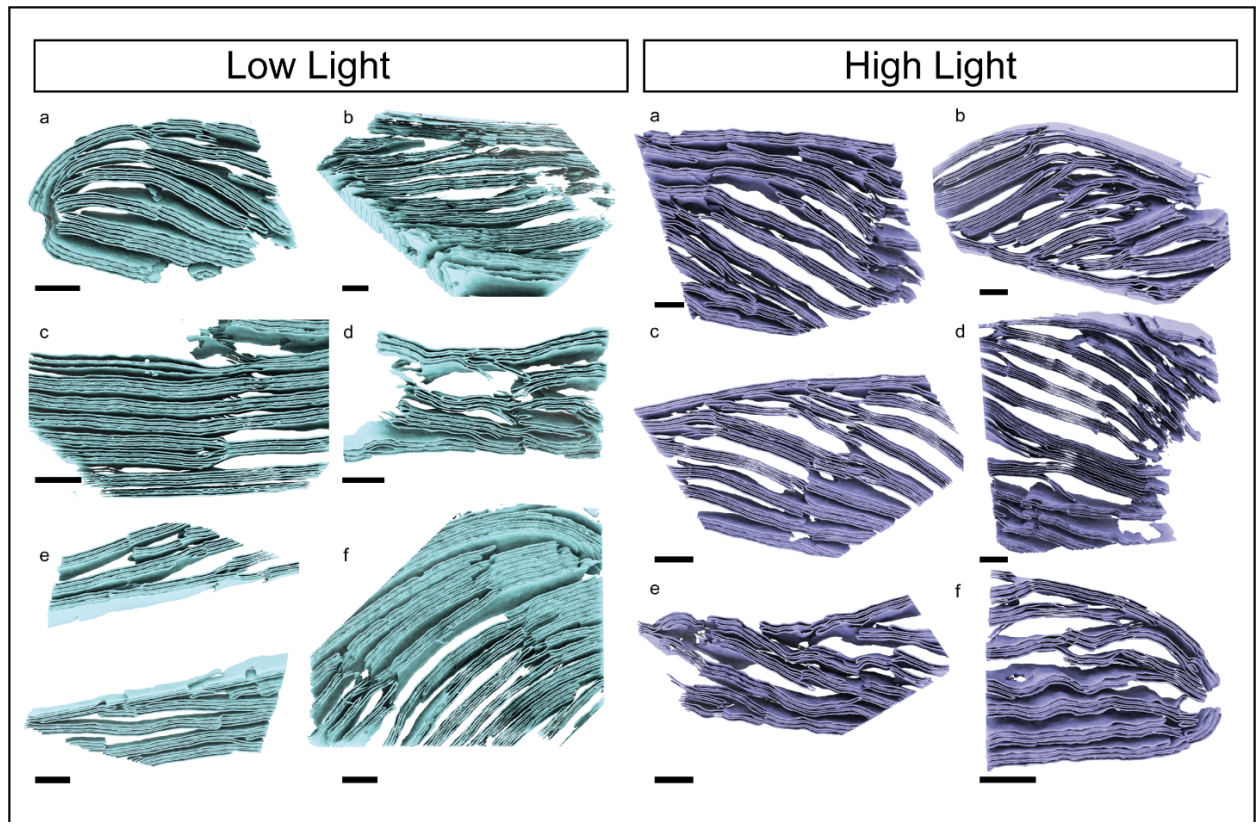

**Figure S4. Gallery of 3D segmentations of thylakoid membranes from *P. tricornutum* cells under LL (left) and HL (right) conditions. Same conditions as in Fig. 3C scale bar 200 $\mu\text{m}$.**

**Supplementary videos are available following the link https://doi.org/10.5281/zenodo.23686172**

LL-movie-tomo04.mp4

LL-movie-tomo01.mp4

HL-tomo02-02.mp4

HL-movie-tomo06.mp4

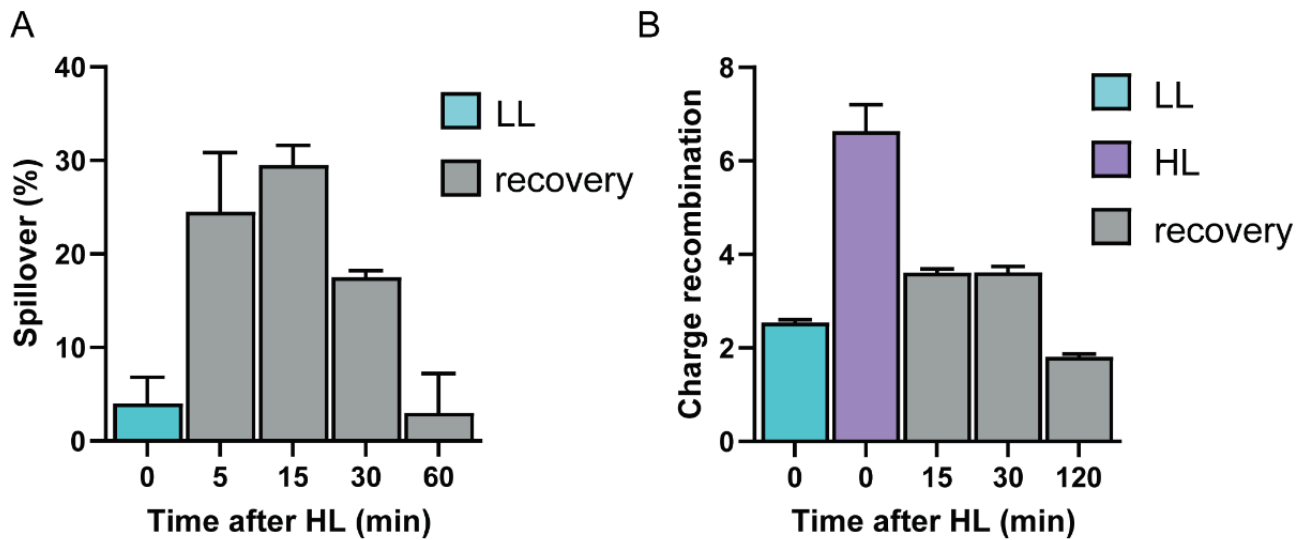

**Figure S5. Recovery kinetics after 2h of HL treatment (A) Spillover, measured as the difference in $P_{700}$ oxidation rate between presence and absence of DCMU. (B) Reversible PSII photoinhibition.**

**Supplementary Table 1** Values of HPLC analysis of the two independent experiments: (diadinoxanthin (Dd); diatoxanthin (Dt); DES: De-epoxidation ratio of the xanthophyll cycle pigments, expressed as the ratio $Dt/(Dt + Dd)$).

|  | FX | FX (d) | DD | DD (d) | DT | DT (d) | DD+DT/FX | DT/ DT + DD |
| --- | --- | --- | --- | --- | --- | --- | --- | --- |
| LL flask1 | 39350380 | 8822441 | 6997225 | 880777 | 595235 | 0 | 0,175892481 | 0,070248832 |
| LL flask2 | 30612836 | 2550675 | 6833421 | 766040 | 406334 | 0 | 0,241403722 | 0,050754984 |
| HL flask1 | 38690736 | 9691438 | 4796391 | 0 | 6298737 | 721982 | 0,244245122 | 0,59411472 |
| HL flask2 | 50054148 | 6625502 | 5298436 | 585777 | 9876805 | 0 | 0,278071901 | 0,626660346 |
| rec flask1 | 30288308 | 8489381 | 8983436 | 93072 | 1626680 | 0 | 0,276014076 | 0,151980886 |
| rec flask2 | 48834312 | 7108769 | 12742997 | 1331043 | 5681611 | 0 | 0,353138416 | 0,287594218 |

| DES | av | sd | *100 | *100 |
| --- | --- | --- | --- | --- |
| LL | 0,060501908 | 0,013784232 | 6,050190803 | 1,378423165 |
| HL | 0,610387533 | 0,023013233 | 61,03875333 | 2,301323308 |
| rec | 0,219787552 | 0,095893106 | 21,97875522 | 9,589310635 |
